# Mapping a brain parasite: occurrence and spatial distribution in fish encephalon

**DOI:** 10.1101/2023.01.26.525646

**Authors:** Ana Born-Torrijos, Gabrielle S. van Beest, Paolo Merella, Giovanni Garippa, Juan Antonio Raga, Francisco E. Montero

**Author notes:** **Corresponding author**: Ana Born-Torrijos.

## Abstract

Parasites, especially brain-encysting trematodes, can have an impact on host behaviour, facilitating the transmission to next host and completion of the life cycle, but insufficient research has been done on whether specific brain regions are targeted. Using the laboratory model *Cardiocephaloides longicollis*, the precise distribution of metacercariae in experimentally-infected, wild and farmed fish was mapped. The brain regions targeted by this parasite were explored, also from a histologic perspective, and potential pathologic effects were evaluated. Experimental infections allowed to reproduce the natural infection intensity of *C. longicollis*, with four times higher infection intensity at the higher dose (150 *vs* 50 cercariae). The observed metacercarial distribution, similar among all fish groups, may reflect a species-specific pattern. Metacercariae occur with highest density in the optic lobe area (primarily infect the periventricular gray zone of optic tectum) and the medulla oblongata, whereas other areas such as the olfactory lobes and cerebellar lobes may be occupied when the more frequently invaded parts of the brain were crowded. Mono- and multicysts (i.e. formed either with a single metacercaria, or with 2 ̶ 25 metacercariae encapsulated together) may be formed depending on the aggregation and timing of metacercariae arrival, with minor host inflammatory response. Larvae of *C. longicollis* colonizing specific brain areas may have an effect on the functions associated with these areas, which are generally related to sensory and motor functions, but are also related to other host fitness traits such as school maintenance or recognition of predators. The detailed information on the extent and distribution of *C. longicollis* in fish encephalon sets the ground to understand the effects of brain parasites on fish, but further investigation to establish if *C. longicollis*, through purely mechanical damage (e.g., occupation, pressure and displacement), has an actual impact on host behaviour remains to be tested under controlled experimental conditions.

## 1 Introduction

Parasites can directly impact host physiology by influencing its growth, reproduction, metabolic or immune functions (e.g. Lochmiller and Deerenberg, 2000; Sánchez et al., 2018) affecting sometimes their behaviour and survival (Poulin, 2013). Some parasites display a number of strategies that enhance their transmission to the next hosts, facilitating the completion of the parasite’s life cycle (Poulin, 2010). Typical direct and indirect phenotypic alterations that parasites have developed to increase their chances of being preyed upon along with their hosts include altered activity, reduced antipredator responses, or changes in host morphology and microhabitat selection (reviewed in Moore, 2002; Lafferty and Shaw, 2013). In fish, brain-encysting parasites have been shown to cause changes in neurotransmitter levels or to damage neural tissue (Shaw and Øverli, 2012), impair shoaling and antipredator behaviour (e.g., Shirakashi and Goater, 2001; Helland-Riise et al., 2020), and to also increase the frequency of conspicuous behaviours that attract bird final hosts, such as flashing, surfacing and contorting (Lafferty and Morris, 1996).

As noted by Helland-Riise et al. (2020), there is insufficient research on whether specific brain regions are targeted by brain-infecting parasites, but this information is necessary to understand specific mechanisms and the neurological processes manipulated by parasites. Parasites that colonise the brain, an immunologically privileged site protected by the blood brain barrier (Cox, 1994), are suitably placed to alter the behaviour of their hosts to enhance their transmission. Just as the patterns of migration to the brain may be species-specific, the site selection or distribution of metacercariae within the brain may also be (Erasmus, 1972). For example, within the Diplostomatidae, the family to which many of the trematodes inducing behaviour changes belong, the species *Posthodiplostomum ptychocheilus* (Faust, 1917) Achatz et al., 2021 (formerly *Ornithodiplostomum ptychocheilus* (Faust, 1917) Dubois, 1936; Achatz et al., 2021) usually occupies the optic lobes and cerebellum of fish (Conn et al., 2008; Hendrickson, 1979), whereas *Diplostomum phoxini* can occupy the optic lobes, but the cerebellum and medulla oblongata are more severely infected (Dezfuli et al., 2007a). Once a trematode larvae, including diplostomatoideans, reach their target organ, they undergo an increase in body size and complexity from the cercarial stage to the infective metacercaria (Matisz et al., 2010a; Conn et al., 2008). The time required to become infective is species-specific, e.g., *Cercariae* X requires 46 days, while *P. ptychocheilus* needs ten weeks to successfully infect the definitive host (Erasmus, 1959; Shirakashi and Goater, 2005). Once metacercariae encyst, they are surrounded by one or two cyst walls that separate the parasite from the host tissue (Hoffman, 1958), but information on the organization of the cyst walls in metacercariae that are encysted individually or closely together with other growing larvae is scarce.

Therefore, to improve the knowledge on brain-encysting trematodes, a species with unknown specific distribution and damage to the brain was selected, the trematode *Cardiocephaloides longicollis* (Rudolphi, 1819) Dubois, 1982 (Strigeidae). This parasite infects the brain of more than 31 fish species from nine families, including numerous sparid species and the farmed *Sparus aurata* L. (Born-Torrijos et al., 2016), and is trophically transmitted to the definitive host, mainly gulls, when infected fish are preyed upon by seabirds. Metacercariae of this parasite have been described to infect the optic lobes of the fish midbrain and, as commonly occurs in brain-encysting trematodes, have been implicated in potential changes to the host’s behaviour including impaired vision and motor control (Prévot, 1974, Recherches sur les cycles biologiques et l’écologie de quelques trématodes nouveaux parasites de *Larus argentatus michaelis* Naumann dans le midi de la France. Université d’Aix-Marseille, France, PhD thesis). Although this hypothesis has been maintained over time (Prévot and Bartoli, 1980; Osset et al., 2005; Bartoli and Boudouresque, 2007; Born-Torrijos et al., 2016), it has never been experimentally tested. In order to prove this, a two-step research is necessary. First characterising the precise distribution of the metacercariae in the different fish brain areas that could justify potential behavioural changes, and second, confirming the potential effects of microhabitat selection in the fish brain with behaviour experiments under controlled conditions. The goal of this study is to accomplish the first step, monitoring the microhabitat selection, encystment and development of the brain-encysting *C. longicollis* in its fish host. To this end, the artificial infection process was first optimized in a laboratory model, using different infection doses. To investigate whether there is a common pattern in the colonization of the different brain areas, the precise distribution of metacercariae was mapped in experimentally-infected as well as in wild and farmed fish, using a histological perspective also to identify the specific targeted regions and their potential pathological effects. With this information, an attempt to link *C. longicollis* distribution to possible effects on host behaviour based on available literature of manipulative trematodes has been done, establishing the base of knowledge for further studies on the actual behavioural changes based on experimental trials to build upon.

## 2 Material and Methods

### 2.1 Host collection and experimental fish infections

Snails *Tritia reticulata* (L.), collected manually in spring at the littoral of an area with a high prevalence of infection (Born-Torrijos et al., 2016; “Beach of the Eucalyptus” (40°37′35.0″N, 0°44′31.0″E) in Els Alfacs Lagoon (Ebro Delta, Spain)), were acclimated and screened for *C. longicollis* infection, by placing them in individual wells for 24 h at 25 °C, 12:12 h light:dark cycle and 35 psu salinity and checking for the emergence of cercariae. The experimental fish were pathogen-free gilthead seabream supplied by a hatchery in Burriana (Castellón, Spain), which were kept in 3000 l tanks at the SCSIE facilities (Central Support Service for Experimental Research, University of Valencia) until used for the experimental infections. Exposure of fish to *C*. *longicollis* cercariae followed the procedure described in van Beest et al. (2019). In brief, freshly emerged cercariae (<2 h old) pooled from different infected snails were placed in a limited volume of water together with fish for 1 hour and 30 minutes. At the end of the experiment, fish were euthanised with an overdose of MS222 (tricaine methanesulfonate, 0.03% solution buffered in seawater; Sigma-Aldrich).

Data on the infection rate of *C. longicollis* metacercariae were obtained from experimental infections using different doses of cercariae (total n=87, see Table 1). To establish a standardized infection protocol and investigate the potential role of infection dose on infection success, experimental fish were infected with either 50 or 150 cercariae (low *vs* high infection dose, respectively), and the number of metacercariae was quantified after one month post-infection (mpi). In an attempt to increase the number of cercariae that an infected fish could survive and to maximize the infection of fish, a potentially lethal dose of 300 cercariae was used in a single infection dose that caused the death of all fish 21 days post infection, whereas control fish that followed the same procedure (with absence of cercariae) and were kept in same conditions suffered no mortality.

**Table 1.** Summary of fish samples used in the present study to investigate various aspects of *Cardiocephaloides longicollis* metacercarial infection, distribution and encystment in the fish brain (see Materials and methods for details). Standard length and weight of fish are given as mean ± SD (min ̶ max). ‘Mean intens.’ indicates mean intensity, ‘Mtc distrib.’ contains the number of samples used for *C. longicollis* metacercariae distribution per brain regions, and ‘Prev’ *C. longicollis* prevalence. Spaces marked in grey indicate that no data were produced for that category.

| Group of fish | No. fish | Standard length (cm) | Weight (g) | Prev (%) | Mean intens. | Mtc. distrib. | Infection rate |
| --- | --- | --- | --- | --- | --- | --- | --- |
| **Experm. infections** |  |  |  |  |  |  |  |
| Lethal dose | 18 | 11.76 ± 1.63<br>(9.50 – 14.50) | 13.33 ± 1.54<br>(10.85 – 15.09) | 100 | 115.8 | – | 38.59±11.83<br>(25.0 – 70.0) |
| Low dose<sup>e</sup> | 33 | 9.20 ± 1.64<br>(5.50 – 13.70) | 23.81 ± 6.10<br>(15.24 – 41.2)<sup>a</sup> | 100 | 7.5 | – | 15.03±13.61<br>(0 – 62) |
| High dose<sup>e,f</sup> | 36 | 13.11 ± 2.03<br>(10.50 – 8.20) | 72.68 ± 55.93<br>(25.8 – 265.1) | 100 | 31.3 | n=14 | 20.89±15.55<br>(4 – 68.67) |
| Both doses | 69 | 11.24 ± 2.70<br>(5.5 – 18.2) | 53.63 ± 49.80<br>(15.24 – 265.1)<sup>b</sup> | 100 | 19.9 | – | 18.09±14.85<br>(0 – 68.67) |
| **Wild fish** |  |  |  |  |  |  |  |
| *S. aurata*<sup>f</sup> | 4 | 29.38 ± 0.95<br>(28.0 – 30.0) | 755.45 ± 118.99<br>(691.6 – 933.8) | 100 | 6.5 | n=4 | – |
| *P. erythrinus* | 10 | 21.75 ± 1.62<br>(20.0 – 25.0) | 270.92 ± 72.54<br>(188.0 – 423.0) | 20 | 3 | n=2 | – |
| *L. mormyrus*<sup>f</sup> | 10 | 21.1 ± 1.05<br>(20.0 – 23.0) | 285.68 ± 17.88<br>(228.0 – 285.78) | 70 | 5.1 | n=7 | – |
| **Farmed *S. aurata*** |  |  |  |  |  |  |  |
| Golfo Aranci (Italy)<sup>f</sup> | 120 | 25.1 ± 1.32<br>(21.0 – 27.0)<sup>c</sup> | 329.01 ± 140.61<br>(115.6 – 582.5) | 40 | 1.3 | n=25 | – |
| Stintino (Italy) | 30 | 24.12 ± 1.24<br>(22.0 – 7.05)<sup>d</sup> | 418.71 ± 16.41<br>(399.0 – 464.0)<sup>d</sup> | 0 | 0 | – | – |
| Marine pond (Spain) | 39 | 24.86 ± 3.1<br>(21.0 – 33.0) | 410.82 ± 61.70<br>(311.0 – 628.86) | 25.6 | 13.1 | half brain<br>n=10 | – |
| Levantine farm (Spain) | 211 | 21.90 ± 2.97<br>(15.5-36.3) | 311.54 ± 143.23<br>(90.79 – 750.4) | 0 | 0 | – | – |
| **Histology** | 9 | 12.29 ± 4.74<br>(8.36 – 22.1) | 95.20 ± 121.00<br>(13.01 – 297.20) | 100 | – | – | – |

<sup>a</sup> based on n=23; <sup>b</sup> based on n=59; <sup>c</sup> based on n=25. Total length is 26.18 ± 4.02 (19.2– 32.7) (n=115); <sup>d</sup> based on n=25; <sup>e</sup> used for statistical analyses testing the effect of infection dose on metacercarial infection rate; <sup>f</sup> used for statistical analyses testing the effect of brain region on No. metacercariae.

### 2.2 Fish dissection and quantification of metacercariae

The number of metacercariae in the brain of different sparid species, including wild and farmed gilthead seabream *S. aurata*, was recorded in a total of 424 fish (Table 1). These included wild-caught specimens of three species of sparids (n=24, *Lithognathus mormyrus* (L.), *Pagellus erythrinus* (L.), *S. aurata* (L.), see Table 1) from commercial fishermen and fish markets and farmed gilthead seabreams from a marine pond (Southern Atlantic, Spain, n=39) and from fish farms in the Eastern and Western Spanish Mediterranean, i.e., from the Levantine coast in Spain (n=211), and two fish farms in Italy (Sardinia, Golfo Aranci, n=120, Stintino, n=30) (Table 1). Of these, a total of 48 naturally-infected fish (including wild and farmed fish, Table 1) were used to investigate the exact distribution of metacercariae in the brain by carefully dissecting different subregions of the brain of infected fish. In addition, the brain of 14 fish infected with a high dose of cercariae were also dissected by subdividing the brain regions.

To investigate whether the parasite has a non-random distribution targeting specific areas of the brain, besides following the descriptions by Wullimann et al. (1996), the different brain regions were subdivided according to the natural division and distinct boundaries of the brain. This was also done to facilitate fine dissection and identification of areas during extensive samplings, when the brain is usually ignored and therefore not analysed. The areas were identified into seven regions (see Figure 1): olfactory bulbs (Olf-B), olfactory lobes (Olf-L), optic lobe regions (Op-L), inferior and superior cerebellar lobes (ICL, SCL), medulla oblongata (Mo), and spinal cord (SC). When possible, the areas were divided by the sagittal plane separating the left and right sides of the brain, but differences in the number of metacercariae between sides were discarded (linear model: log(No. metacercariae+1) ~ Brain side + No. total metacercariae, estimate=−0.0076, SE=0.0197, p=0.699), and data from both sides were pooled. Eyes were also analysed to exclude infections, as described for other *Cardiocephaloides* spp. (Timi et al., 1999; Vermaak et al., 2021). All fish were measured and weighed prior to dissection, and experimental fish were also measured before experimental infection to assess whether the infection rate was related to size. All brain region dissections and quantification of metacercariae were performed under a stereomicroscope (Leica MZ APO, Germany, equipped with camera Leica DFC295).

**Figure 1.**
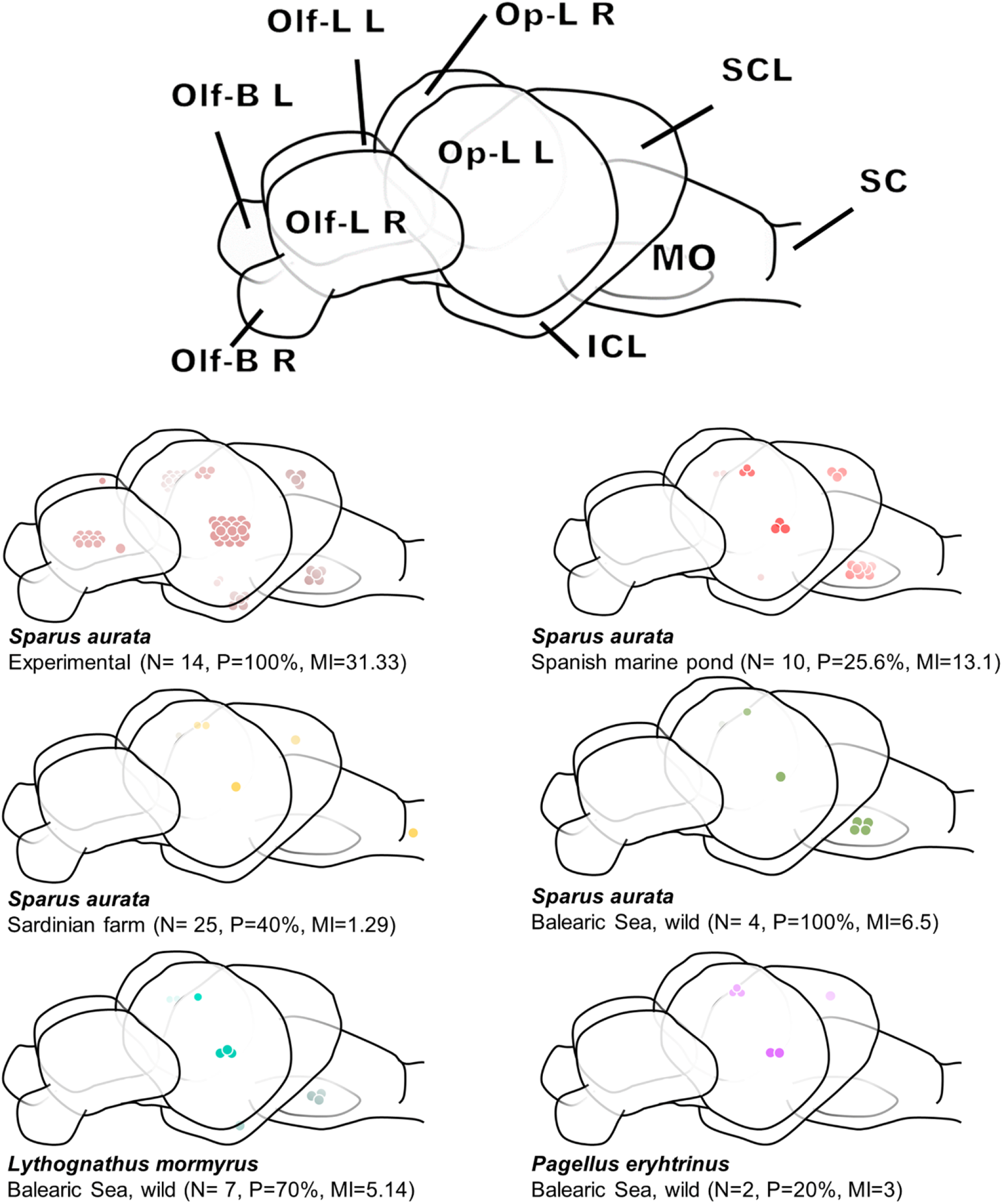
Distribution of metacercariae of *Cardiocephaoides longicollis* in the different fish brain regions, i.e., olfactory bulbs (Olf-B), olfactory lobes (Olf-L), optic lobe regions (Op-L), inferior and superior cerebellar lobes (ICL, SCL), medulla oblongata (Mo), and spinal cord (SC). Metacercarial distribution in different fish species sampling locations are provided. N, number of infected brains used for metacercarial distribution; P, prevalence (based on total number of fish, see Table 1); MI, mean intensity. Note that the number of metacercariae in the brain of fish from the marine pond is based only of half brain (see Materials and methods).

### 2.3 Statistical analyses: infection dose and distribution in the brain

To examine whether there was an effect of infection dose on the number of encysted metacercariae in the fish brain, i.e., the infection rate, a linear model was fitted, using ‘number of metacercariae’ as the response variable, and ‘infection dose’ (low *vs* high, i.e., 50 *vs* 150 cercariae) and ‘fish size’ (standard length, mm) as covariables.

To investigate possible differences in the number of metacercariae per brain region between different fish groups, two linear models (LMs) were fitted using ‘number of cysts/region’ as the response variable (log transformed), and ‘brain regions’ (Olf-B, Olf-L, Op-L, ICL, SCL, Mo, SC) and ‘fish group’ as well as their interaction as fixed variables, with the number of total cysts in the brain as a covariable to control for different total cyst numbers. The fish groups tested were (i) experimentally-infected *vs* wild gilthead seabream (*S. aurata*) or (ii) wild gilthead seabream *vs* striped seabream (*L. mormyrus*). Other comparisons could not be statistically done as the number of fish was too small or only partial data were available (i.e. only half of the brains of gilthead seabream from the marine pond were dissected because the other half were used for histology, but the samples were not properly fixed and therefore could not be used, Table 1), but were included in the figures to visually represent the distribution of metacercariae. Data were statistically analysed using R (R Core Team, version 4.1.2), with the significance level set at 0.05, and the R^2^ values of the models performed were reported (*MuMIn* package, Bartón, 2022).

### 2.4 Ultrastructure and histology of metacercariae in fish brain

The formation of the cyst from metacercariae after reaching the target tissue was studied in nine experimentally-infected fish (Table 1), infected with 100 or 150 cercariae and dissected for histological analyses at different post-infection times (6, 21 days post infections (dpi), and 8 and 15 months post infection (mpi)). The brain and eyes were fixed in 10% buffered formalin for two days, dehydrated in graded alcohol and embedded in paraffin. Paraffin blocks were sectioned (5-7 μm, 20-40 sections per block) using a microtome (Leica RM 2125RT, Germany). Histological sections were stained with hematoxylin and eosin, mounted in DPX, and examined at 100 x to 400 x magnification (Leica DMR, Germany, equipped with a Leica DMC5400 camera).

To investigate the cysts formed by *C. longicollis* metacercariae in the brain, their ultrastructure was examined in isolated cysts collected from experimentally-infected fish (8 mpi) and wild fish (*S. aurata* and *L. mormyrus*). In addition, metacercariae encysted in brain tissue were collected from wild fish (*S. aurata*). All samples were fixed in 2.5% glutaraldehyde in phosphate-buffered saline (PBS, 0.1 mol L^−1^, pH 7.4) and preserved at 4 °C for 48 h. Samples were then washed three times in PBS (15 min each), post-fixed in 2% osmium tetroxide (2 h), washed three times in phosphate buffer (PB), and dehydrated in a graded series of acetone. Samples were then infiltrated and embedded in Epon resin. First, semithin sections of 400 nm were cut using a glass knife, stained with toluidine blue, and observed under a light microscope. Then, ultrathin sections (60–90 nm) were cut through selected regions using a diamond knife on a Leica Ultracut UCT ultramicrotome. These were placed on copper grids and stained with uranyl acetate and lead citrate according to Reynolds (1963). For observation a JEOL JEM-1010 transmission electron microscope was used (JEOL, Tokyo, Japan, accelerating voltage 80 kV, equipped with a CCD digital camera Mega View III at the Laboratory of Electron Microscopy, Institute of Parasitology, Biology Centre of the Czech Academy of Sciences (Czech Republic)).

## 3 Results

### 3.1 Quantification of *C. longicollis* metacercariae, experimental infection dose and infection rate

Metacercariae of *C. longicollis* were found in all groups of fish, with variable infection levels among experimentally-infected, wild and farmed fish (Table 1). In the farmed fish group, *C. longicollis* was found only in one of the Italian fish farms with a prevalence of 40% while it was absent from the Spanish Levantine coast. The low sample size in wild fish is due to these specimens being obtained for the analysis of metacercarial distribution in the brain (see below) rather than to study the prevalence of *C. longicollis* in these species, which is reported in Born-Torrijos et al. (2016).

All fish used in the infection experiments were successfully infected with a prevalence of 100% and varying infection intensity depending on the fish group. The mean intensity of infection was higher in the experimentally-infected fish (7.5 ̶ 115.8) than in the wild (3 ̶ 6.5) and the farmed ones (1.3 ̶ 13.1). No *C. longicollis* were found in the eyes of any of the fish. The extremely high intensity observed in the fish infected with 300 cercariae in a single dose (115.8 mean metacercariae/fish, 75 ̶ 210 min-max metacercariae) was likely the reason why the fish did not survive longer than 21 dpi, since all 18 fish died within two days (while 25 uninfected fish from the same fish lot kept in the same conditions suffered no mortality). Therefore, a dose of 300 cercariae was considered lethal in fish from 10 to 15 cm. The infection dose showed a positive effect on the number of metacercariae that successfully arrived in the fish brain, meaning that using a higher dose significantly increased the infection rate of *C. longicollis* metacercariae (LM, estimate=0.0107, SE=0.0028, p<0.001), with no effect of fish size (LM, estimate=0.1024, SE=0.0522, p=0.054) (Suppl. Table S2, Figure S1).

### 3.2 Metacercariae distribution in the fish brain

Metacercariae were found in all the seven regions of the fish brain, except for the olfactory bulb (the most anterior region, Fig. 1) and the spinal cord (the most posterior; except for one metacercaria in *S. aurata* from a Sardinian farm). The most heavily infected fish were the experimental gilthead seabream, however, metacercarial distribution was only slightly different depending on the fish origin and species (Figs. 1 and 2). While experimentally-infected fish showed a clear increase in metacercariae in the Op-L region, wild and farmed *S. aurata* from the marine pond showed a higher number of metacercariae encysted in the Mo, as did wild *L. mormyrus* and *P. erythrinus*. Comparisons between experimentally-infected and wild gilthead seabreams showed certain differences in the number of metacercariae in different brain regions. While in experimentally-infected fish, the Op-L showed the highest number of metacercariae compared to the Olf-B (LM, estimate=3.444, SE=0.2663, p<0.001), followed by the Olf-L, ICL and Mo (Suppl. Table S1), the number of metacercariae in wild *S. aurata* was comparable between regions, with a more pronounced difference in the Op-L region in experimentally-infected fish.

**Figure 2.**
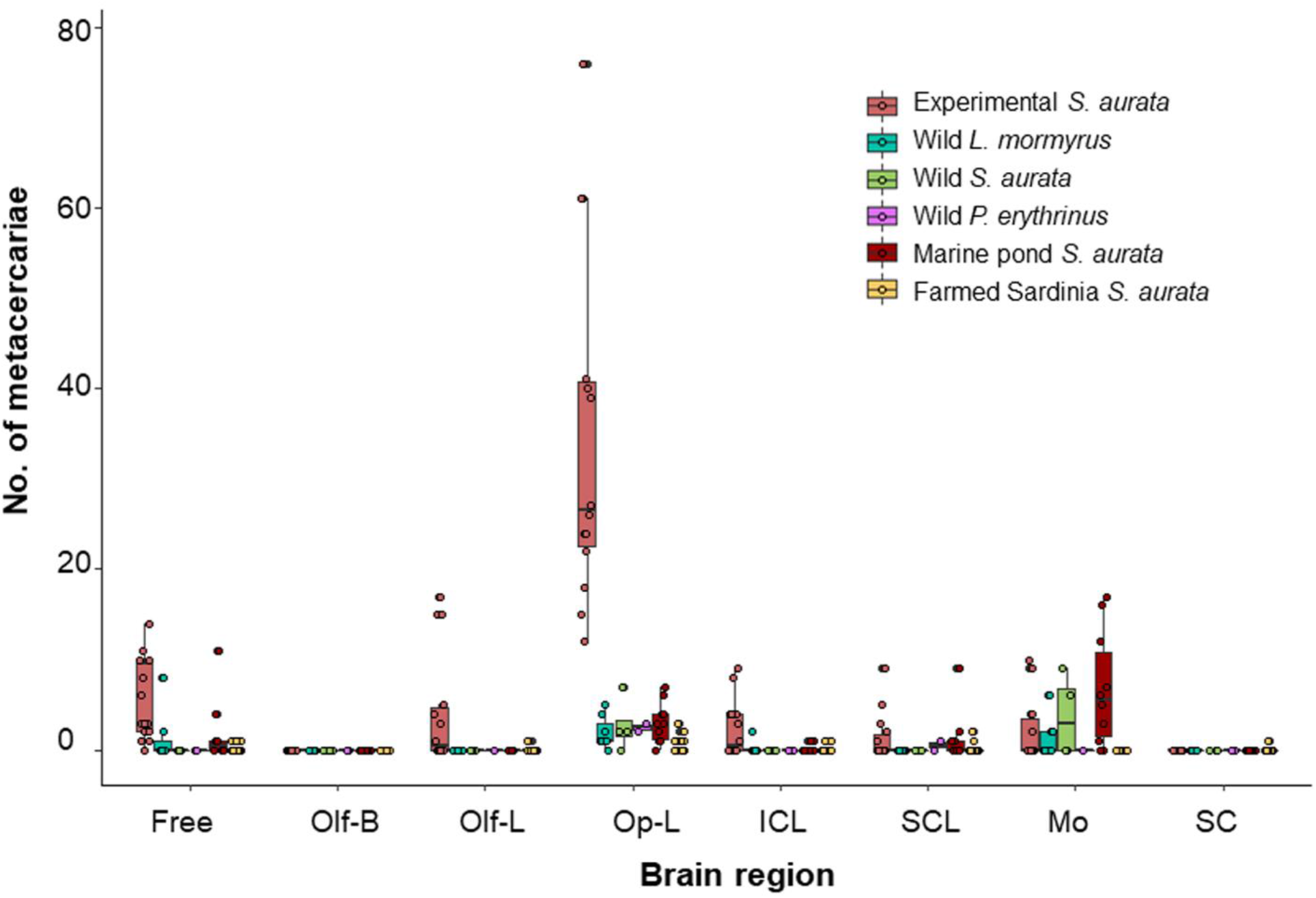
Variation in the number of metacercariae of *Cardiocephaloides longicollis* encysted in different fish groups. Box plots represent the median number of metacercariae per brain region, upper and lower quartile (box) with maximum and minimum ranges (whiskers) and outliers (grey circles). Dots represent jittered raw data.

The number of metacercariae in the brain regions of both wild gilthead and striped seabream was similarly distributed, being significantly higher in the Op-L and Mo regions than in the Olf-B in the striped seabream (estimate=0.940, SE=0.2481, p<0.001; estimate=0.5919, SE=0. 2481, p=0.020, Suppl. Table S1).

To describe the distribution of metacercariae, the Op-L region was included as an integrated area as certain structures such as the periventricular gray zone of optic tectum (PGZ) represent a continuum that cannot be to cleanly separated during dissection. In freshly dissected brains, the presence of metacercariae, including multicysts (see later; Figs. 3 and 4), was observed within the PGZ and bellow the medulla oblongata (Fig. 3, A, B), but to identify the specific location, histology was required to distinguish the different regions according to cell types. When the cercariae arrived in the brain, they were found at 6 dpi occupying only the tectal ventricle (Fig. 3C) with no occupation within the surrounding tissues. When metacercariae grew (21 dpi, 8 and 15 mpi) they occupied a larger part of the ventricle and also migrated to the PGZ, but not to the tectum opticum (TeO, part of the Op-L, see Fig. 3 D). There were no remarkable pathologies associated with the metacercariae presence revealing that even though some of the *C. longicollis* metacercariae accumulate in the tectal ventricle, most did not occupy the optic lobe itself but were located in the PGZ and on the cerebellum (Fig. 3 E, F). In any case, the tectum opticum (TeO) was not occupied in any of the slides, with metacercariae being found in the PGZ when the optic lobe was infected.

**Figure 3.**
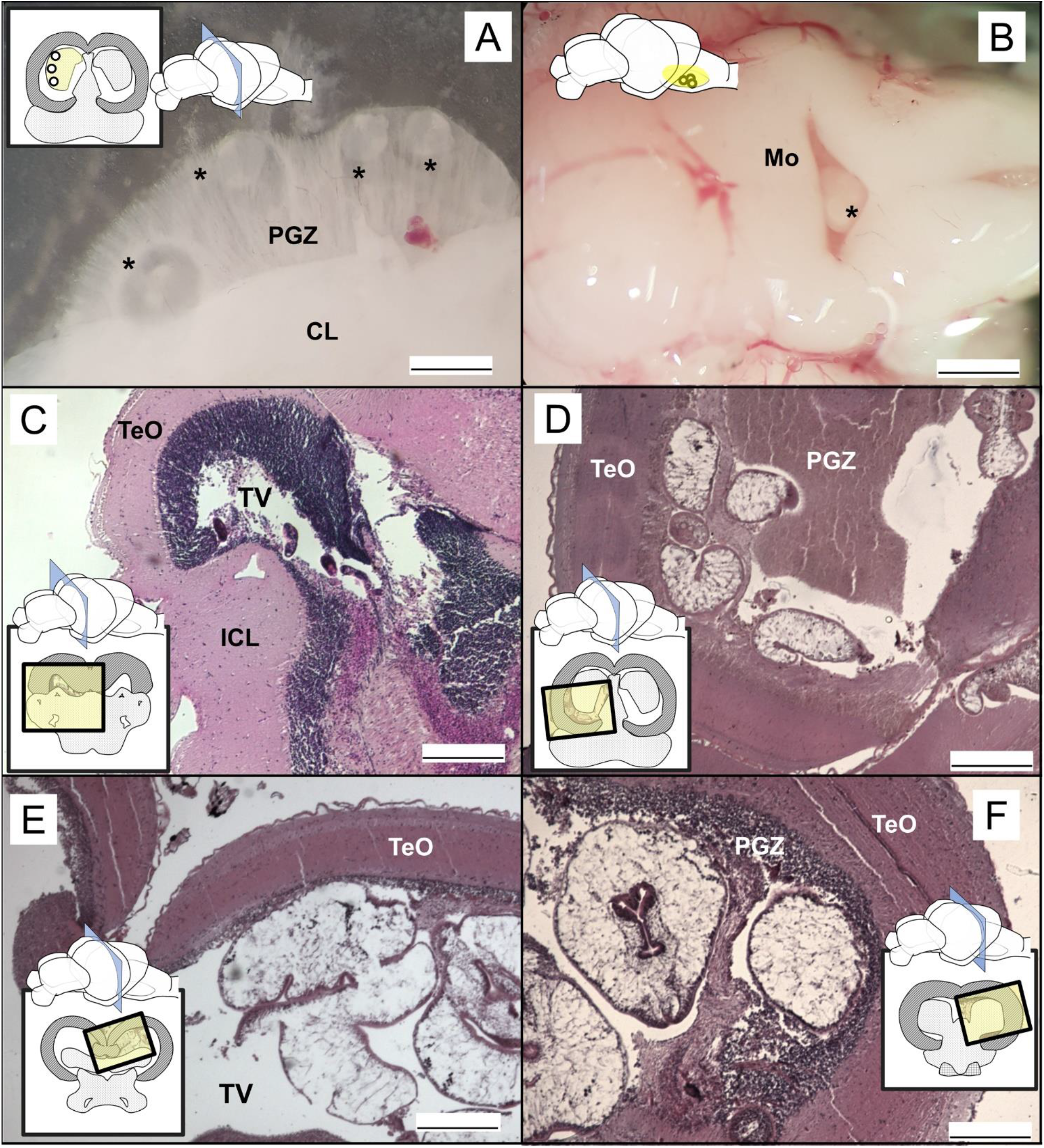
Occupation of the fish brain by *Cardiocephaloides longicollis* in fresh (A, B) and histological samples (C-F). *Cardiocephaloides longicollis* metacercariae within (A) the PGZ and (B) the medulla oblongata in experimentally-infected fish one month after infection. Asterisks indicate the position of metacercariae. *Cardiocephaloides longicollis* metacercariae are found at 6 dpi in the tectal ventricle (C), and as they grow (D, 21 dpi; E, 8 mpi; F, 15 mpi) they occupy larger part of the tectal ventricle, and also the PGZ. The representations of brains indicate the sections and positions (yellow square) where metacercariae have been found. Legend: TeO striped, cerebellum in dots and Mo squared. ICL, inferior cerebellar lobe; Mo, medulla oblongata; PGZ, periventricular gray zone of optic tectum; TeO, tectum opticum; TV, tectal ventricle. Scale bars: A= 300 μm; B = 450 μm; C-F= 200 μm.

**Figure 4.**
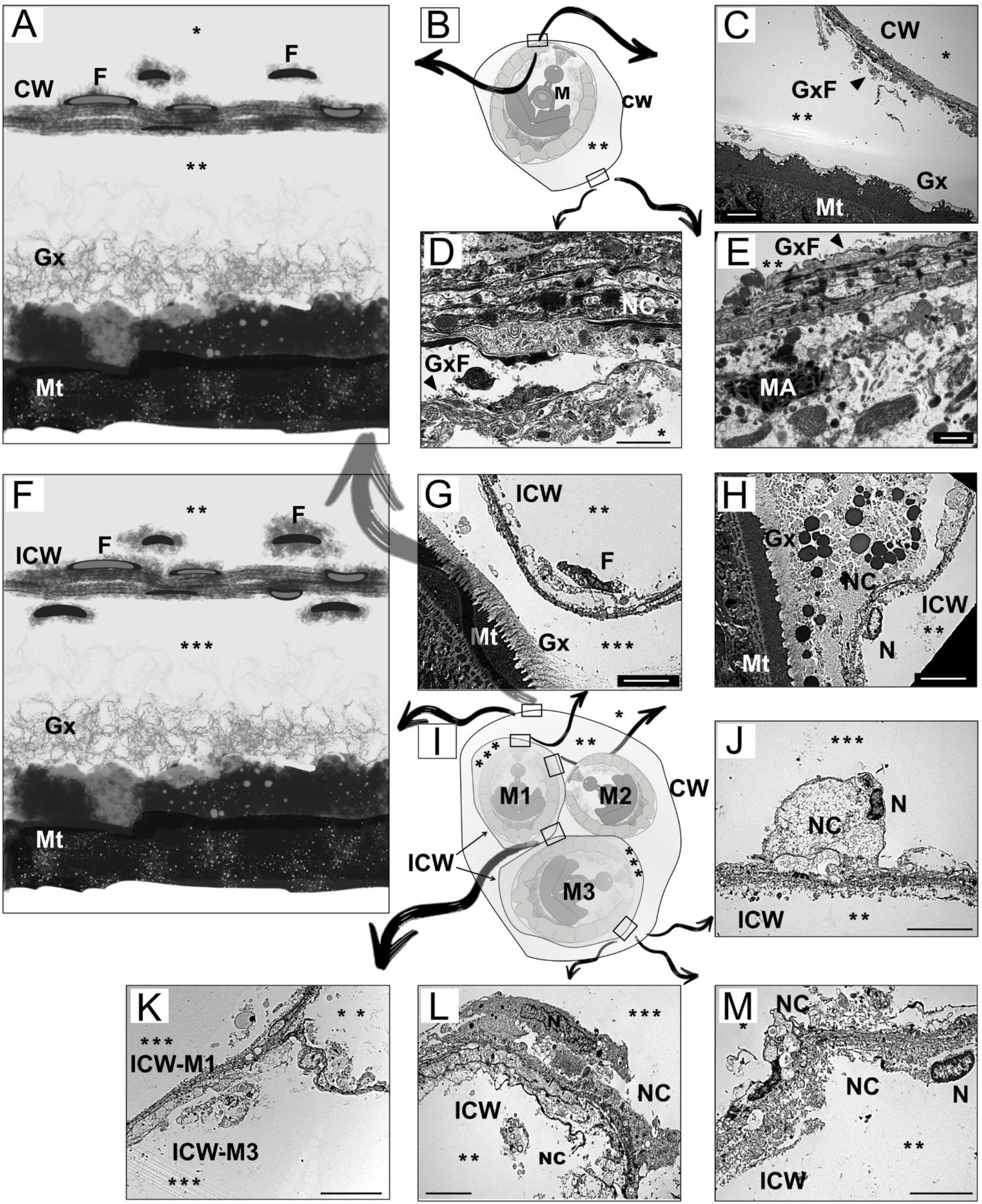
Transmission electron micrographs showing the tegument and capsule walls of metacercariae of *Cardiocephaloides longicollis*. A and F illustrate the capsule wall of monocyst and multicyst metacercariae; B and I represent diagrams of monocyst and multicyst metacercariae showing the location of the following TEM micrographs. C – E Longitudinal section through the capsule wall and tegument of a monocyst. G, H, J-M Longitudinal section through the inner capsule wall and tegument of a multicyst metacercaria. D, E, J-M Detail of necrotic material accumulated on the capsule wall surrounding the metacercaria. K, Detail of inner capsule walls merging together within a multicyst. CW, capsule wall; F, fibrocyte; Gx, glycocalyx; GxF, glycocalyx filaments; ICW, inner capsule wall; M, metacercaria; M1-M3 number of metacercaria in a multicyst; MA, macrophage; Mt, metacercarial tegument; N, nucleus; NC, necrotic cells. Head arrows indicate glycocalyx filaments, asterisks (*) outside of the cyst, (**) inside of the cyst, *** inside of the cyst when encysted with more than one capsule wall. Scale bars: d, E =1 μm; C, G, H, J, L, M = 5 μm; F = 10 μm.

### 3.3 Mono- and multicysts of *C. longicollis*

Cercariae of *C. longicollis* were found in the brain by 6 dpi with a similar morphology to that of free-swimming cercariae (Fig. 3). However, by 21 dpi the parasite increased considerably in size and the tegumental structures changed to resemble metacercariae rather than free-living cercariae, but lacking a visible cyst. Larvae found at 8 and 15 mpi had the normal morphology of encysted metacercariae that had developed a cyst in the brain. These were surrounded by a wall, and filled with a viscous glycocalyx-like material that maintained the position of the parasite within the cyst. Nevertheless, the host surface showed little disruption in response to the parasite.

Two types of cysts of *C. longicollis* metacercariae were observed in the brain of captured and experimentally-infected fish: (i) a monocyst in which a single metacercaria was wrapped by a capsule (Fig. 4 B), or (2) a multicyst in which 2 to 25 metacercariae were encapsulated together (Fig. 4 I). Whereas the first type had a single wall in direct contact with the host, the multicyst may have a common capsule surrounding all metacercariae, or each metacercariae may produce its own capsule as a single parasite, likely resulting in metacercariae with more than one capsule. The composition of the capsule walls, both inner and outer, was identified by TEM and was composed mostly of host cells in necrosis and filamentous material possibly originated from the glycocalyx between the tegument of the parasite and the capsule (Fig. 4 C, G). The capsule was produced by the parasite so that the material accumulated in the inner layers of the capsule wall (Fig. 4 C-D), where the necrotic cells were probably produced by the host tissue when it came into contact with the parasite. Thus, the necrotic cells appeared to accumulate on the outer layers of the capsule of monocysts and also in the outermost capsule of a multicyst (Fig. 4 A-E). However, the composition of the wall in each individual capsule within a multicyst was different, showing cells in necrosis on both sides of the wall (Fig. 4 F-M). In this case, the degraded material was not only in the outer, but also in the inner layers of the capsule wall, resulting in a thick but loose capsule wall (Fig. 4 L, M) that likely merged with other inner wall capsules close to it (Fig. 4 K). Although the immune response was likely to be very low, TEM revealed that the capsule wall was characterised by the presence of fibrocytes that were flattened in the inner layers of the wall. In addition, the host tissue in contact with the cyst showed the occasional presence of macrophages (Fig. 4 E).

## 4 Discussion

The optimised experimental infection protocol allowed for the reproduction of the natural infection intensity of *C. longicollis* and showed that metacercariae occur with the highest intensity in the optic lobe area. Higher resolution examination of the brain revealed that the cercariae occupy the PGZ area as early as six days after infection and remain with no occupation within the surrounding tissue, growing into larger encysted metacercariae that may be single or grouped in multiple cysts. The higher intensity in the optic lobe area and the medulla oblongata may point to a targeted site selection occupying the brain areas encoding motor, visual and basal physiological functions. However, directly connecting the occupation of these areas to effective behavioural changes remains to be confirmed with behaviour experiments.

As expected, the different doses used resulted in different infection rates, with more than four times higher intensity of infection at the higher dose. Although an infection intensity of ~116 metacercariae was achieved with the highest dose used (300 cercariae), the concentration of larvae was too high and became lethal to ~12 cm long fish. Prévot and Bartoli (1980) reported hyperinfection with *C. longicollis* and subsequent death of two other sparid species, but did not provide information on infection dose and intensity. The highest intensity recorded in wild fish hosts is ~67 metacercariae in another sparid species (*Diplodus annularis*, 150 ̶ 162 mm SL, Born-Torrijos et al., 2016, Suppl. Table S7), so it is not surprising that this dose was lethal. The lethality observed a few days after infection could also be related to the simultaneous arrival and disruption of brain tissue when cercariae begin to grow. Cranial distortion has been described to occur following the simultaneous arrival of cercariae in the brain, as many metacercariae develop in a time interval that exceeds the host’s ability to accommodate for them, interfering with normal development, and likely contributing to brain tissue destruction and reduced fish survival (Sandland and Goater, 2001).

The presence of *C. longicollis* is reported in farmed gilthead seabream from a new aquaculture facility in Italy, with relatively low prevalence compared to that found in a Spanish fish farm by Born-Torrijos et al. (2016). However, the negative data from Spanish farms on the Levantine coast suggest that the occurrence of the parasite in aquaculture net pens may have a patchy distribution, likely related to several factors such as seafloor topography, ocean currents, the presence of suitable hosts in the surrounding, and even the ability of larvae to reach their next host (van Beest et al., 2022). Fish from brackish water ponds showed a higher infection intensity than other farmed fish likely because these ponds are in natural spaces and have an influx of parasites from the wild surrounding fauna through the external water supply they typically require (Bouwmeester et al., 2021). The transmission of *C. longicollis* to fish could thus be enhanced if intermediate hosts are concentrated in these ponds with slower water renewal, but further snail sampling would be needed to confirm this. The absence of *C. longicollis* in the eyes of any experimentally-infected, wild and farmed fish (see also Born-Torrijos et al., 2016) is noteworthy because other *Cardiocephaloides* spp. and various brain-encysting larvae have also been found in the eyes (Hendrickson, 1979; Timi et al., 1999; Muzzall and Kilroy, 2007; Vermaak et al., 2021).

The symmetrical distribution of *C. longicollis* metacercariae in both cerebral hemispheres seems to be a common feature (Siegmund et al., 1997). Nevertheless, the metacercariae followed a non-random distribution in the brain of the experimentally-infected, wild and farmed hosts, with a similar pattern and only minor exceptions in certain areas of experimentally-infected and wild specimens (Fig. 1) with this site selection likely reflecting a species-specific pattern in metacercarial distribution. There were no significant differences in the distribution of metacercariae between wild striped seabream and wild gilthead seabream, whereas experimentally-infected gilthead seabream had a higher concentration of metacercariae in the optic lobe area than the wild gilthead seabream. This could be related to the higher overall intensity in experimentally-infected fish as other areas also had higher infection rates (19.94 *vs* 6.5, Figs. 1 and 2).

The olfactory bulb (Olf-B) was the only area completely free of metacercariae of *C. longicollis*, and only a single metacercaria was found in the spinal cord of a farmed fish from Italy. Dezfuli et al. (2007a) showed in another diplostomatoidean, *D. phoxini*, the absence of metacercariae in the olfactory bulb, as well as the olfactory lobes, the inferior lobe of the cerebellum and the pituitary. However, Barber and Crompton (1997) suggested that this parasite species, infecting in this case a different fish species, could occupy the olfactory lobes and the spinal cord when the more frequently invaded parts of the brain were crowded. This might not be the case with *C. longicollis* as the spinal cord was free of infection even in experimentally heavily infected fish. However, the medulla oblongata, the region closest to the spinal cord (Fig. 1), had a higher presence of *C. longicollis* metacercariae in wild and farmed fish except in specimens from Golfo Aranci, which also had a lower overall intensity. It cannot be thus ruled out that cercariae occupy areas near their entry point, as suggested in species aggregating in brain regions after likely following the optic and cranial nerves (Hendrickson, 1979; Helland-Riise et al., 2020). Although cercariae of *C. longicollis* have been observed penetrating areas likely connected to migration routes to neuronal canals and also detected in the connective tissue of the oral cavity of the fish 4 hours post infection (van Beest et al., 2019, 2022), the exact route of migrating cercariae remains unknown. They could also use the muscle or circulatory system to reach the target organ, as do other diplostomatoidean (e.g. Erasmus, 1959; Ratanarat-Brockelman, 1974; Hendrickson, 1979; Haas et al., 2007; Matisz et al., 2010b; Matisz and Goater, 2010).

With all this together, it can be suggested that the primary site of arrival for *C. longicollis* might be the optic lobe area, more specifically the PGZ, and the medulla oblongata, whereas other areas such as Olf-L and ICL/SCL are occupied depending on the intensity of infection. This would also explain the higher occupation of the latter areas in the experimentally-infected fish, as metacercariae would accumulate there when the optic lobe and medulla oblongata are occupied by the sudden arrival of a high number of metacercariae. The “preference” of *C. longicollis* for the optic lobe area and medulla oblongata seems to be consistent with other brain-encysting parasites including diplostomatids (Hendrickson, 1979; Radabaugh, 1980a; Barber and Crompton, 1997; Dezfuli et al., 2007; Helland-Riise et al., 2020).

Metacercariae are usually considered immature, non-feeding and resting stages (Bush et al., 2001), but prior to cyst formation and this resting phase, diplostomatoidean larvae that infect the brain normally undergo a variable and extended period of development before encystment (Erasmus, 1972; Dönges, 1969; Conn et al., 2008). Larvae of *C. longicollis* are described, for the first time, occupying the tectal ventricle and specifically the PGZ cavity as early as 6 dpi, increasing their size considerably, from ~100 to ~400 μm within three weeks, with no cyst formation during this period (Fig. 3), a similar growth to that described by Prévot and Bartoli (1980) (this refers to size increase; note that the present measurements refer to fixed and histological material and the publication referred to fresh larvae). Metacercariae require at least four months of development before becoming infective to their final host (Prévot and Bartoli, 1980), likely reaching their maximum growth rate during this time as do other diplostomatoideans (e.g. Sandland and Goater, 2000; Shirakashi and Goater, 2005). In some species, newly arrived larvae might change their position over time, as described for diplostomatid metacercariae, i.e. after reaching the brain and initially dispersing in deeper areas of the brain, larvae appear to move and concentrate in the outermost layers of the optic lobes and cerebellum after a few days of infection (Conn et al., 2008; Matisz et al., 2010a; Matisz and Goater, 2010). This has not been observed in our data as *C. longicollis* remains occupying a larger section of the PGZ area until at least 15 months post infection (Fig. 3F), not moving to other regions and even not infecting the tectum opticum, thus likely only generating mechanical pressure on it. Therefore, in the optic lobes area *C. longicollis* appears to specifically occupy the PGZ area, an empty space below the lobes, rather than “sitting” on them as is the case in some species (see Plate 2 in Hoffman and Hoyme, 1958, and Figure 6 in Matisz et al., 2010b).

Microlamellar and microvillar teguments have been described in developing metacercariae to allow their growth, expanding and forming a net around the larva (Goater et al., 2005). The tegumental filaments observed in *C. longicollis* could be the remnants of microvilli as these disappear at early stages (ca. 28 dpi, Goater et al., 2005). The filaments were surrounded by a glycocalyx-like material, as described for *P. ptychocheilus* (Goater et al., 2005), occupying the space between the parasite and the cyst wall. In addition, the presence of two cyst walls has been described, with inner and outer cysts, similar to other trematodes including diplostomatids (Hoffman, 1958; Halton and Johnston, 1982; So and Wittrock, 1982). The second or further cysts are likely to be formed when a new metacercaria arrives to an area already occupied by another larva and the two are then encapsulated together forming a double cyst. Because cyst walls in the brain are usually thin compared with cysts in other tissues (Halton and Johnston, 1982), the inner and outer layers are sometimes not easily distinguished. Monocysts of *C. longicollis* had a very thin inner layer composed largely of loose filaments of the parasite tegument surrounded by a thicker layer composed mostly of host cells, e.g., fibrocytes, with the degree of degradation varying depending on when these cells where incorporated into the outer layer. A similar interface of necrotic material/cell degeneration has been described between the inner and outer cyst layers (Mitchell, 1974; Halton and Johnston, 1982). Similarly, when *C. longicollis* is surrounded by two cysts, the outer cyst had host cells in the outer part and a thin inner layer in contact with the inner cyst had a thicker and lax cyst wall with disorganised structure. This phenomenon is probably due to host cells surrounding a metacercarial cyst that became entrapped in a new cyst, probably caused by the presence of a metacercaria nearby. Fibrocytes and macrophages are observed in the cyst areas where the host tissue and parasite cyst are in direct contact (Fig. 4 E), concentrated near the PGZ. Fibroblasts in the vicinity of the cyst have also been described (So and Wittrock, 1982; Matisz et al., 2010b) as a classic host inflammatory response to the parasite cyst wall or its excretory products (Halton and Johnston, 1982). Other than that, no severe pathological damage to brain tissue was observed, including the presence of rodlet cells which are related to the host inflammatory defence response and have been described in brain-encysting trematodes (Dezfuli et al., 2007; Matisz et al., 2010b). As Dezfuli et al. (2007b) noted for *Diplostomum* sp., the host cellular reaction has been largely overlooked in histopathological studies with the rodlet cells mostly absent from the literature until recently. Further TEM studies of metacercariae in the brain may help to identify the presence of these cells. Some studies have described massive inflammation of the meninges caused by encysted metacercariae (Matisz et al., 2010a) and even large cleared areas with a low proportion of the nervous tissue remaining, surrounded by cellular debris in heavily-infected fish (Dezfuli et al., 2007). However, brain-infecting parasites generally do not appear to cause severe inflammation or pathology in brain tissue (this study; Halton and Johnston, 1982; Siegmund et al., 1997; Goater et al., 2005), with the most common effects being changes in host behaviour. Alterations in host visual acuity have been associated with developing stages rather than with the encysted stages (Shirakashi and Goater, 2005), possibly related to direct penetration of parasite microvillar filaments into the host intercellular matrix or due to the feeding habits of developing metacercariae, which has only been demonstrated in two diplostomatids (Bibby and Rees, 1971; Goater et al., 2005). Further studies on *C. longicollis* are necessary to investigate whether this is an extended process.

Prévot (1974, Recherches sur les cycles biologiques et l’écologie de quelques trématodes nouveaux parasites de *Larus argentatus michaelis* Naumann dans le midi de la France. Université d’Aix-Marseille, France, PhD thesis) suggested that *C. longicollis* may provoke changes in fish behaviour due to impaired vision and motor control of swimming that would cause body oscillations and make fish flanks more visible and thus more prone to be preyed upon by the final hosts, seabirds. Nevertheless, to confirm that a parasite is causing these changes, specific behaviour experiments need to be performed. The present data are therefore not enough to support the existence of these changes in fish hosts infected with *C. longicollis*. However, in order to plan these time-consuming behavioural experiments, evidence that the parasite is occupying areas that could potentially lead to behaviour changes is necessary, and this is the ultimate aim of the present study. Therefore, the functions of the areas occupied by *C. longicollis* have been evaluated in relation to the potential impact that this parasite may have in fish hosts. Learning, memory and avoidance behaviours, as well as regulation of locomotion and basal physiology have been attributed to areas in the forebrain and cerebellum of fish (Wullimann et al., 1996; Rodríguez et al., 2005; O’Connell and Hofmann, 2011). These areas, including the olfactory lobes and superior and inferior cerebellar lobes, were occupied mostly in experimentally-infected fish. However, the primary infection sites across all fish were in the optic lobe region, specifically the PGZ, and in the medulla oblongata. In general, metacercariae colonizing the optic lobe region and medulla oblongata are capable of altering the sensory and motor functions in fish (Barber and Crompton, 1997; Dezfuli et al., 2007a). The function of the medulla oblongata has also been linked with involuntary responses such as breathing and digestion (Bernstein, 1970; Wullimann et al., 1996).

Moreover, the optic tectum is recognized as the primary vision-processing centre in teleosts, responsible for receiving and integrating visual stimuli, including movement, color and shape, and also playing a role in integrating information received from the lateral line system (Springer et al., 1977; Weale, 1982; Guthrie, 1986; Wullimann et al., 1996). The midbrain, which includes the optic tectum, also regulates reproductive physiology (Wullimann et al., 1996). This broadens the traits that could be affected, as damage to the optic tectum is clearly associated with several host fitness traits such as school maintenance, prey detection or recognition of predators, as has been described in fish infected with other diplostomatoideans (Hendrickson, 1979; Radabaugh, 1980b; So and Wittrock, 1982; Barber and Crompton, 1997; Sandland and Goater, 2001). Other parasites have been described occupying the innermost layer of the meninx (Matisz et al., 2010a), whose main function in fish is the secretion of cerebrospinal fluid and maintenance of the blood-brain barrier (Schwarz et al., 1993; Hoffmann and Schwarz, 1996). The “free” metacercariae in the cavity could have resulted from the detachment of these layers during manipulation of the brain, but no detachment was observed in histological samples and their presence could be natural in fluid-filled cavities as described for *P. ptychocheilus* (e.g., Radabaugh, 1980b).

Colonization of the nervous system by *C. longicollis* could by mere mechanical damage (i.e., occupation, pressure, traction or displacement of the tissue) impair the swimming ability or visual acuity of their hosts even if not destroying the tissue or causing severe physical pathologies. Whether this occupation of specific brain regions directly causes behavioural changes is something that needs to be tested under experimental conditions with designs that include transcriptomic and proteomic analyses that can provide further insight into the mechanisms involved in host behavioural changes.

## Acknowledgments

We acknowledge the core facility Laboratory of Electron Microscopy (LEM), Biology Centre of ASCR institution, supported by the MEYS CR (LM2018129 Czech-BioImaging) and Central Service for Experimental Research of the University of Valencia (Spain), specifically the microscopy service and the experimental aquarium plant. We thank the Andromeda Group (Spain) for kindly providing the fish used in this study, and Clemente Graziano (Compagnie Ittiche Riunite and Emanuele Depalmas (Palma d’Oro) (Italy) for their collaboration in fish sampling. The authors are grateful to Phillip Riekenberg for improving the English text.

## Author Contributions

A.B.-T., F.E.M. and G.S.B. conceived the ideas and designed the methodology; A.B.-T. and G.S.B. performed the experimental work, with help of P.M. for fish dissection and of F.E.M. for analysis of histological material; A.B.-T. and F.E.M. obtained the ethics approval; A.B.-T. supervised the project and acquired the funding, did the formal data analysis and wrote the first draft of the manuscript. All the authors contributed critically to the draft and gave final approval for the publication.

## Data Availability Statement

The dataset underlying this article will be archived in FigShare repository upon acceptance of the manuscript.

## Ethics statements

This study was approved by the Ethics Committee of the University of Valencia (Spain), authorisation procedure A1390575271192 (2014/037/UVEG/001, Dirección General de Agricultura, Ganadería y Pesca, Valencian Local Government).

## Supplementary Materials

**Supplementary Table S1**. Results of linear model (LM) evaluating the differences in number of metacercariae of *Cardiocephaloides longicollis* depending on the brain region and the group of fish ((i), experimental *vs* wild gilthead seabream, and (ii) striped seabream *vs* wild gilthead seabream) (LM, log(No. metacercariae) ~ Brain region × Fish group + No. total metacercariae). The intercept contains the reference variable ‘Olf-B’ and (i) ‘experimental gilthead seabream’ or (ii) ‘wild striped seabream’. R^2^ represents the variability explained by fixed effects, and statistically significant results are in bold (α=0.05).

**Supplementary Table S2**. Results of linear model (LM) evaluating the differences in number of encysted metacercariae of *Cardiocephaloides longicollis* depending on infection dose (low *vs* high), using the fish standard length as covariable (LM, No metacercariae ~ Fish size + Infection dose). The intercept contains the reference variable low infection dose. R^2^ represents the variability explained by fixed effects, and statistically significant results are in bold (α=0.05).

**Figure S1**. Variation in the number of encysted metacercariae of *Cardiocephaloides longicollis* depending on the infection dose. Box plots represent the median number of metacercariae per infection dose, upper and lower quartile (box) with maximum and minimum ranges (whiskers) and outliers (grey circles). Dots represent jittered raw data.

