## Supplementary for "Mapping a brain parasite: occurrence and spatial distribution in fish encephalon"

*** Corresponding author:**

Ana Born-Torrijos

**Table S1**. Results of linear model (LM) evaluating the differences in number of metacercariae of *Cardiocephaloides longicollis* depending on the brain region and the group of fish ((i), experimental *vs* wild gilthead seabream, and (ii) striped seabream *vs* wild gilthead seabream) (LM, log(No. metacercariae) ~ Brain region × Fish group + No. total metacercariae). The intercept contains the reference variable 'Olf-B' and (i) 'experimental gilthead seabream' or (ii) 'wild striped seabream'. $R^2$ represents the variability explained by fixed effects, and statistically significant results are in bold (α=0.05).

| | LM | Estimate | SE | t-value | P |
|---|---|---|---|---|---|
| (i) | $R^2$= 0.69 | | | | |
| | Intercept (Olf-B; experim. fish) | -0.4078 | 0.2332 | -1.749 | 0.083 |
| | **Olf-L** | **0.9940** | **0.2663** | **3.732** | **<0.001** |
| | **Op-L** | **3.444** | **0.2663** | **12.932** | **<0.001** |
| | **ICL** | **0.8148** | **0.2663** | **3.059** | **0.003** |
| | SCL | 0.5195 | 0.2663 | 1.950 | 0.054 |
| | **Mo** | **0.6441** | **0.2663** | **2.418** | **0.017** |
| | SC | $1.126^{e-15}$ | 0.2663 | 0.000 | 1.000 |
| | wild *S. aurata* | 0.3550 | 0.4171 | 0.851 | 0.397 |
| | **TotalBrain** | $\mathbf{8.133^{e-03}}$ | $\mathbf{2.743^{e-03}}$ | **2.965** | **0.004** |
| | OlfL_total × wild *S. aurata* | -0.9940 | 0.5650 | -1.759 | 0.081 |
| | **OpL_total** × wild *S. aurata* | **-2.3750** | **0.5650** | **-4.204** | **<0.001** |
| | ICL_total × wild *S. aurata* | -0.8148 | 0.5650 | -1.442 | 0.152 |
| | SCL_total × wild *S. aurata* | -0.5195 | 0.5650 | -0.919 | 0.360 |
| | Mo_total × wild *S. aurata* | 0.4180 | 0.5650 | 0.740 | 0.461 |
| | SC_total × wild *S. aurata* | $1.670^{e-16}$ | 0.5650 | 0.000 | 1.000 |
| (ii) | $R^2$= 0.44 | | | | |
| | Intercept (Olf-B; wild str. seabream) | -0.1304 | 0.1982 | -0.658 | 0.513 |
| | Olf-L | $1.934^{e-15}$ | 0.2481 | 0.000 | 1.000 |
| | **Op-L** | **0.940** | **0.2481** | **3.788** | **<0.001** |
| | ICL | 0.1569 | 0.2481 | 0.633 | 0.529 |
| | SCL | $9.663^{e-16}$ | 0.2481 | 0.000 | 1.000 |
| | **Mo** | **0.5919** | **0.2481** | **2.386** | **0.020** |
| | SC | $9.801^{e-16}$ | 0.2481 | 0.000 | 1.000 |
| | wild *S. aurata* | 0.0344 | 0.2919 | -0.118 | 0.907 |
| | TotalBrain | 0.0254 | 0.0179 | 1.415 | 0.162 |
| | OlfL_total × wild *S. aurata* | $-9.489^{e-16}$ | 0.4114 | 0.000 | 1.000 |
| | OpL_total × wild *S. aurata* | 0.1293 | 0.4114 | 0.314 | 0.754 |
| | ICL_total × wild *S. aurata* | -0.1569 | 0.4114 | -0.381 | 0.704 |
| | SCL_total × wild *S. aurata* | $-8.437^{e-16}$ | 0.4114 | 0.000 | 1.000 |
| | Mo_total × wild *S. aurata* | 0.4702 | 0.4114 | 1.143 | 0.257 |
| | SC_total × wild *S. aurata* | $-9.226^{e-16}$ | 0.4114 | 0.000 | 1.000 |

**Table S2.** Results of linear model (LM) evaluating the differences in number of encysted metacercariae of *Cardiocephaloides longicollis* depending on infection dose (low *vs* high), using the fish standard length as covariable (LM, No metacercariae ~ Fish size + Infection dose). The intercept contains the reference variable low infection dose. $R^2$ represents the variability explained by fixed effects, and statistically significant results are in bold (α=0.05).

| LM | Estimate | SE | t-value | P |
|---|---|---|---|---|
| $R^2$= 0.48 | | | | |
| Intercept (low dose) | 0.3098 | 0.4358 | 0.711 | 0.480 |
| Standard length | 0.1024 | 0.0522 | 1.962 | 0.054 |
| **High dose** | **0.0107** | **0.0028** | **3.841** | **<0.001** |

**Figure S1.** Variation in the number of encysted metacercariae of *Cardiocephaloides longicollis* depending on the infection dose. Box plots represent the median number of metacercariae per infection dose, upper and lower quartile (box) with maximum and minimum ranges (whiskers) and outliers (grey circles). Dots represent jittered raw data.

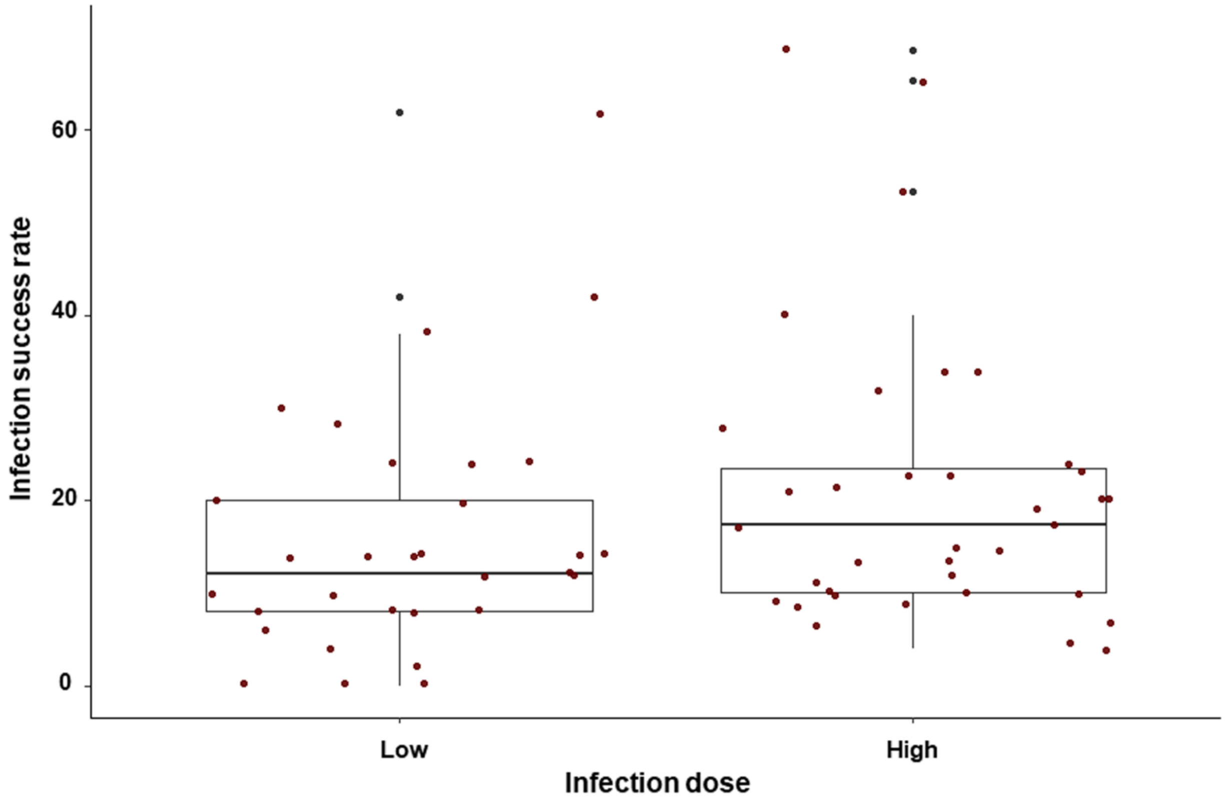


*Supplementary data in this article will be archived in Figshare repository upon acceptance of the manuscript.*
